# Where conservation and restoration matter most: sensitivity-based prioritization of connected forest

**DOI:** 10.64898/2026.09.10.750632

**Authors:** Bram Van Moorter, Rannveig M. Jacobsen, K. Ulrika J. Asplund, Yennie K. Bredin, Jenni Nordén, Graciela Rusch, Bálint Czúcz, Manuela Panzacchi

**Affiliations:** Norwegian Institute for Nature Research (NINA), Sognsveien 68, 0855 Oslo, Norway

## Abstract

Achieving global conservation and restoration targets requires knowing not only how much land to conserve and restore, but also where such actions will have the greatest ecological effect. Conservation and, in particular, restoration priorities are often inferred from habitat condition, opportunity, or broad ecological value, yet these criteria do not necessarily identify where local changes generate the largest landscape-scale consequences. Building on sensitivity-based conservation prioritization, we extend the approach to derive ecological loss and gain potential by differentiating a persistence-relevant connectivity metric with respect to local habitat condition. Sensitivity identifies areas where local degradation or improvement would most strongly affect connected habitat at the landscape scale. Combining sensitivity with current condition distinguishes high loss potential, where degradation would have large ecological consequences, from high gain potential, where ecological improvement could generate large benefits. Applying the framework to national forest-naturalness data in Norway, we show that loss and gain potential are spatially related but far from redundant, and that gain potential is not simply concentrated in the most degraded forests. Both high-loss and high-gain areas were poorly represented within current protected areas, although loss potential was consistently better represented than gain potential. Extending sensitivity-based prioritization in this way provides a general and scalable means of linking local habitat-condition change to persistence-relevant landscape outcomes, while distinguishing the ecological potentials that can inform conservation and restoration decisions.

## Introduction

Biodiversity loss has reached a scale that demands stronger protection of remaining high-quality ecosystems (IPBES 2019). Protection alone, however, will not be sufficient to reverse current declines, and ecological restoration is therefore becoming an equally important component of biodiversity policy (Mace et al. 2018; Díaz et al. 2020; Banks-Leite et al. 2020). The Kunming–Montreal Global Biodiversity Framework commits countries to conserve 30% of terrestrial, inland-water, coastal, and marine areas by 2030 and to restore 30% of degraded ecosystems (CBD 2022), while the UN Decade on Ecosystem Restoration (UNEP & FAO 2021) and the EU Nature Restoration Regulation (European Parliament and Council of the European Union 2024) further reinforce restoration as a central component of biodiversity policy. Meeting these ambitions will require major investments in land, management, and restoration, yet land is contested and both conservation and restoration are costly (Strassburg et al. 2020). Spatial prioritization is therefore unavoidable: limited resources should be directed toward places where protection prevents the greatest ecological losses and restoration generates the greatest ecological gains. The central challenge is not only how much land to conserve or restore, but also where action will contribute most to biodiversity persistence.

The ecological consequences of changing habitat condition depend not only on how much suitable habitat remains, but also on how that habitat is distributed and connected across the landscape (Suding 2011; Gilby et al. 2018). Habitat loss and degradation reduce the amount and quality of habitat available locally, while fragmentation alters the spatial relationships among habitat areas and can restrict movement, thereby limiting migration, dispersal, colonization, and rescue processes that underpin population persistence (Hanski 1999; Fahrig 2003; Haddad et al. 2015; Nordén et al. 2013). These processes interact. The ecological importance of any one location therefore depends not only on its local condition, but also on the amount and quality of surrounding habitat and on how readily organisms can move among habitat areas (Beger et al. 2022). Protecting a suitable site or improving a degraded one within a well-connected habitat network may make a disproportionate contribution to biodiversity persistence, whereas an equivalent action at an isolated site may have much smaller effects (Hunter-Ayad & Hassall 2020; Li et al. 2023). Conservation and restoration planning should therefore account not only for local habitat condition, but also for how changes in that condition alter habitat availability and connectivity across the wider landscape (Daigle et al. 2020; Beger et al. 2022; Frey et al. 2012; Li et al. 2023).

Spatial prioritization has a long history in conservation, where systematic conservation planning provides a formal framework for identifying sets of areas that best achieve conservation objectives under spatial and resource constraints (Margules & Pressey 2000; Moilanen et al. 2009). Across conservation and restoration, many prioritization approaches – particularly in restoration planning – evaluate candidate sites primarily from local attributes such as ecological condition, threat, degradation, feasibility, costs, or expected benefits, using expert judgement, ranking, scoring, or cost–benefit criteria (Noss et al. 2009; Orsi & Geneletti 2010; Brancalion et al. 2019; Strassburg et al. 2020). Landscape-ecological and green-infrastructure approaches extend this perspective by incorporating spatial context, for example through habitat amount, proximity, neighbourhood structure, or connectivity, and have been used both to identify areas important for maintaining ecological networks and to locate opportunities for strengthening them through restoration (Daigle et al. 2020; Hanson et al. 2022; Tambosi et al. 2014; Ortega et al. 2023; Bubnicki et al. 2024; Wang et al. 2025). These approaches improve the representation of landscape structure, but typically use landscape context as a property of candidate sites rather than directly estimating how changing habitat condition at a particular location would alter landscape-scale habitat availability, connectivity, and ultimately biodiversity persistence.

One way to move beyond static descriptions of habitat condition and landscape context is to explicitly evaluate how changing a particular spatial unit affects a landscape-level ecological outcome. Such perturbation-based prioritization has been developed through both explicit and analytical approaches. Explicit approaches evaluate the effect of removing, adding, or modifying spatial units by recomputing landscape connectivity after each change. They can accommodate relatively detailed representations of movement through heterogeneous landscapes, including least-cost and random-walk or circuit-based formulations, and have been applied to both conservation and restoration analyses (Saura & Torné 2009; García-Feced et al. 2011; McRae et al. 2012; Foltête et al. 2014). Their main limitation is computational because the landscape model must be rerun for every candidate perturbation. Analytical approaches, by contrast, derive the effect of habitat change directly from the structure of the landscape model. In metapopulation theory, for example, eigenvector–eigenvalue relations were used to quantify how habitat destruction, restoration, and the contribution of individual habitat patches affect landscape-level metapopulation persistence (Ovaskainen 2003; Ovaskainen & Hanski 2003). These approaches avoid repeated perturbation but have generally been formulated for patch-based landscape representations in which movement through the intervening landscape is implicit. Moreover, eigenanalysis itself becomes increasingly demanding as the number of habitat units grows. High-resolution conservation and restoration planning therefore requires an approach that combines the spatial realism of explicit movement models with the computational advantages of analytical sensitivity.

Our approach combines these two traditions. We use spatially explicit connectivity models that represent movement through heterogeneous landscapes, but summarize habitat configuration and connectivity using a landscape-scale measure of habitat functionality (hereafter, ‘landscape functionality’), such as metapopulation capacity (Hanski & Ovaskainen 2000) or probability of connectivity (Saura & Pascual-Hortal 2007). Following the analytical logic of Ovaskainen (2003) and Ovaskainen and Hanski (2003), Van Moorter et al. (2026) derived a sensitivity framework for such landscape-functionality measures across connectivity models representing movement strategies along a continuum from optimal to random movement (see also Kivimäki et al. 2024). The framework allows the effects of local changes in habitat quality or permeability to be quantified without repeated model evaluation. Here, we build on that framework by treating habitat condition as a common driver of both habitat quality and permeability and using its sensitivity to quantify ecological change in both directions: losses through degradation and gains through restoration.

The sensitivity analysis quantifies how strongly landscape functionality responds to a small change in habitat condition at a particular location. Where sensitivity is high, deterioration of habitat condition can cause a disproportionate loss in landscape functionality, whereas improvement can generate a disproportionate gain. We therefore distinguish loss potential, representing the ecological consequence of degrading existing habitat condition, from gain potential, representing the ecological consequence of improving it. These quantities contribute to conservation and restoration priority, respectively, but do not constitute complete management priorities. Applied prioritization may additionally depend on factors such as threat, restoration feasibility, costs, land ownership, and governance. Here, we focus on the ecological component of that broader decision problem by identifying where degradation or restoration is expected to have the greatest consequences for persistence-relevant landscape functionality.

We apply this framework to forest landscapes in Norway, where the ecological consequences of conservation and restoration depend strongly on both local habitat condition and spatial configuration. In boreal and boreonemoral forests, many species of conservation concern are associated with old or near-natural stands, large trees, dead wood, structural complexity, long temporal continuity, and large or spatially connected areas of suitable forest (Majdanová et al. 2023; Jacobsen et al. 2020; Asplund et al. 2024; Esseen et al. 1997; Siitonen 2001; Jonsson et al. 2005, Artsdatabanken 2021). These include saproxylic beetles, wood-inhabiting fungi, polypores, lichens, and bryophytes. At the same time, intensive forestry has reduced both the abundance of old-growth structures and the spatial and temporal continuity of near-natural forest across Fennoscandian landscapes (Östlund et al. 1997; Bubnicki et al. 2024; Wang et al. 2025). Spruce-, pine-, and broadleaf-dominated forests support partly distinct species communities (Tikkanen et al. 2006; Jonsell et al. 2007; Hägglund & Hjältén 2018; Burner et al. 2021) and differ in conservation status, restoration potential, and connectivity challenges (Wang et al. 2025). We therefore represent separate ecological networks associated with spruce-, pine-, and broadleaf-dominated forests and evaluate them under contrasting movement scales. Using these networks, we ask whether sensitivity-based analysis can (1) identify where changes in forest condition have the greatest effects on landscape functionality; (2) determine whether the effects of forest-condition change arise primarily through local habitat quality or through landscape permeability; (3) reveal spatial differences between ecological loss and gain potential; and (4) assess how areas of high loss and gain potential are represented within current protected areas.

## Materials and Methods

### Study area, system and ecological assumptions

We applied the framework to forest landscapes in mainland Norway at a spatial resolution of 50 × 50 m. The study area spans a broad environmental gradient, including boreal and boreonemoral forests, deciduous mountain forests, and smaller areas of lowland nemoral forest. Most forest area is boreal or boreonemoral and varies substantially in tree-species composition, management history, and degree of naturalness (Fig. 1). The national raster comprised 131,544,774 spatial units. Landscape functionality and sensitivity to local changes in forest condition were evaluated across the national raster using local moving-window calculations, allowing national-scale analysis while retaining spatially explicit representations of habitat configuration and movement. See Figure 2 for the analytical workflow.

**Figure 1.**
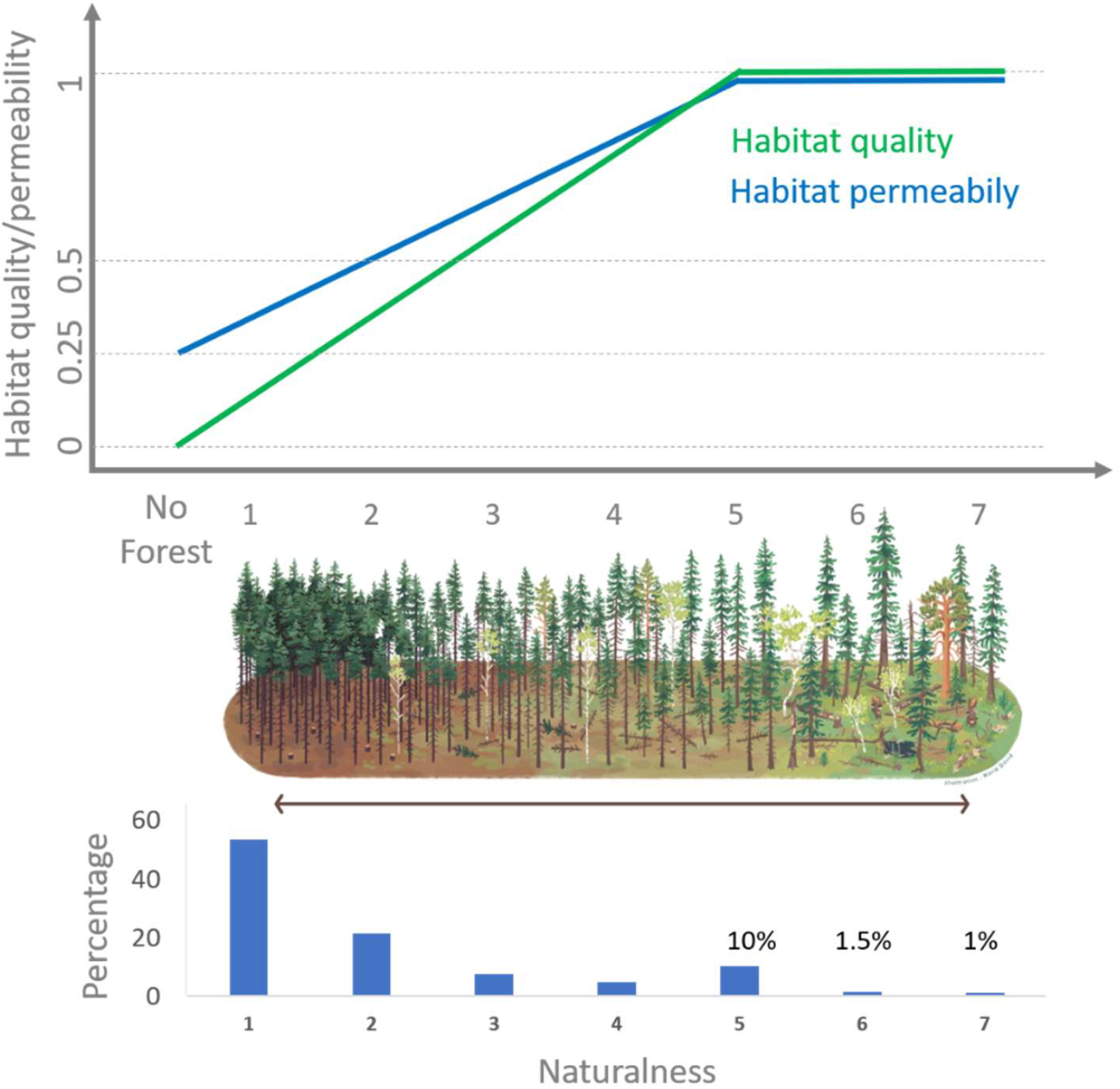
Conceptual representation of forest condition or naturalness and its relationship to habitat quality and habitat permeability. Forest stands are grouped into seven naturalness classes ranging from lowest naturalness (1) to highly natural forest (7). Habitat quality (green) and permeability (blue) increase linearly with naturalness and plateau at class 5; non-forest areas have quality 0, whereas their permeability (i.e. as matrix) is 0.25. The central illustration provides a conceptual visualization of the structural gradient from intensively modified to more natural forest, and the lower panel shows the distribution of forest area among naturalness classes in Norway (classes 6 & 7 cover less than 3% of the landscape).

**Figure 2.**
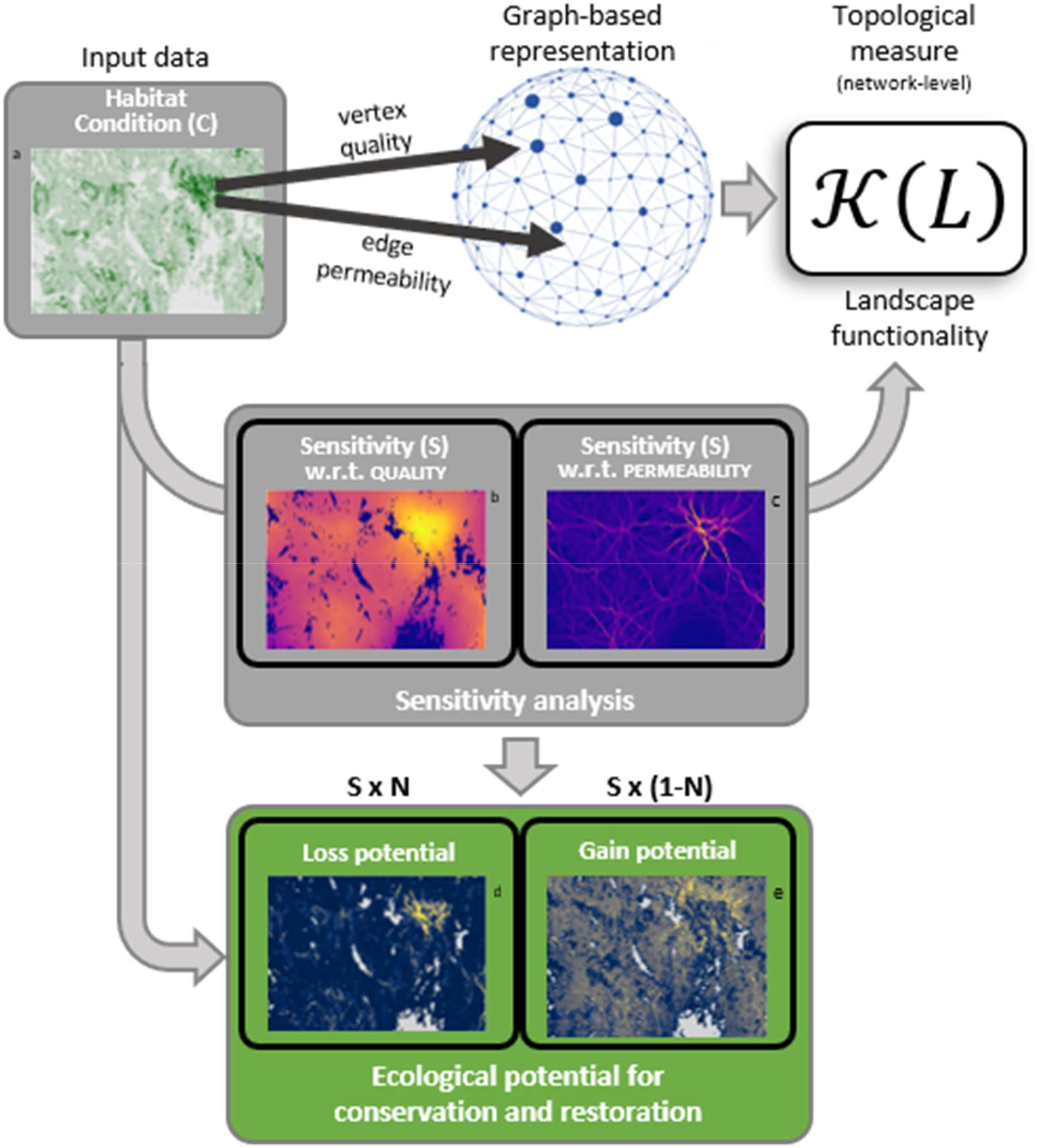
Conceptual workflow of the sensitivity-based prioritization framework. Local habitat condition determines both the habitat quality q_i_ of spatial unit i and its contribution to landscape permeability. A connectivity model is then used to estimate pairwise ecological proximity k_st_ between spatial units s and t, from which the landscape matrix m_st_ = q_s_q_t_k_st_ is constructed. Landscape functionality, K(L), summarizes this matrix to quantify the amount and connectedness of habitat in landscape L. Sensitivity measures how a small local change in habitat condition affects landscape functionality through two pathways: changes in habitat quality and changes in landscape permeability. Combining sensitivity with current habitat condition yields ecological loss potential, representing the potential consequence of degradation, and ecological gain potential, representing the potential benefit of improvement. These ecological potentials can inform, but do not by themselves determine, conservation and restoration priorities.

The analysis targets biodiversity associated with natural and near-natural boreal and boreonemoral forests rather than any single focal species. Forest biodiversity depends not only on forest cover, but also on structural and compositional characteristics that determine habitat quality, including tree species, old and large trees, dead wood of different dimensions and decay stages, microhabitats, microclimatic conditions, and temporal forest continuity (Betts et al. 2022; Matricardi et al. 2020; Fritz et al. 2008; Majdanová et al. 2023; Jacobsen et al. 2020; Asplund et al. 2026). These habitat requirements occur across several taxonomic groups, including saproxylic beetles, wood-inhabiting fungi, polypores, lichens, and bryophytes (Lachat et al. 2025).

We represented forest-associated biodiversity using three broad forest habitat groups associated with spruce-, pine-, and broadleaf-dominated forests (Tikkanen et al. 2006; Jonsell et al. 2007; Hägglund & Hjältén 2018; Burner et al. 2021), crossed with two movement scales to yield six landscape networks. The movement scales were chosen to span dispersal distances reported as relevant for forest-associated insects, fungi, and other small forest organisms (Ranius et al. 2011; Sverdrup-Thygeson et al. 2014; Komonen & Müller 2018). Empirical dispersal estimates for a single species can vary substantially among studies and methodologies. For example, for *Phlebia centrifuga*, Norros et al. (2012) concluded from direct measurements that effective colonization may become dispersal-limited within tens to a few hundred metres, whereas Edman et al. (2004) detected relationships with surrounding old spruce forest at kilometre scales. Assigning species unambiguously to fixed dispersal classes was therefore not feasible for this study. The six networks should instead be interpreted as plausible combinations of forest association and movement scale rather than as models parameterized for particular species (Lachat et al. 2025; Lapin et al. 2025).

### Spatial data and representation of forest condition

We characterized each 50 × 50 m spatial unit using spatial data on forest extent, tree-species composition, and forest condition. All input layers were harmonized to a common 50 × 50 m grid aligned with the Statistics Norway grid. All source datasets are publicly available.

Forest extent was derived from the ‘National base map for area analysis’ (Nasjonalt grunnkart for arealanalyse). We selected areas classified as “forest and woodlands” and rasterized their coverage to the 50 × 50 m analysis grid. Forest presence was subsequently treated as binary, such that any cell containing forest was assigned a value of 1 and cells without forest a value of 0. Areas outside the forest mask were not assigned habitat quality, although they remained part of the movement matrix through the permeability model.

Tree-species composition was derived from the ‘SR16 forest resource raster map’ (Skogressurskart 16 × 16 m raster – SR16R; Breidenbach et al. 2021; Astrup et al. 2019). We used the dominant-tree-species layer, which distinguishes spruce, pine, and broadleaf forest, and resampled it to estimate the proportion *T*_*i,g*_ of each 50 × 50 m cell *i* associated with ecological group *g*. Cells with mixed tree-species composition could therefore contribute habitat to more than one ecological network. The broadleaf category combines boreal and temperate broadleaved tree species because these are not separated in the source product.

Forest condition, (*C*_*i*_), was represented using the ‘similarity-to-natural-forest’ layer (naturskogsnærhet) from the Norwegian natural-forest maps (Hauglin et al. 2025). The source layer expresses forest naturalness on a seven-class scale. We merged classes 5−7 because the two highest classes were extremely rare (Fig. 1), yielding a five-class scale that was rescaled to 0.2–1. For each 50 × 50 m analysis cell, forest condition was calculated as the mean of the underlying 16 × 16 m values.

### Habitat quality and permeability

For each ecological group, *g*, we translated local forest composition and forest condition into two spatially explicit model quantities: ‘habitat quality’, which determines how strongly a cell contributes as habitat, and ‘habitat permeability’, which determines how readily movement can occur through that cell (Fig. 1).

Habitat quality of spatial unit *i* for ecological group *g* was defined as:

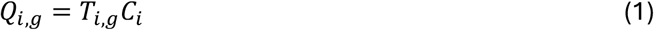

where *T*_*i,g*_ is the proportion of the cell associated with forest type *g*, and *C*_*i*_ is forest condition. Habitat quality therefore increases jointly with the amount of the relevant forest type and its condition, and is zero where the focal forest type is absent.

Habitat permeability was defined across the entire landscape. Non-forest cells were assigned a baseline permeability (*B* = 0.25), reflecting the assumption that movement through the matrix is more difficult but not impossible. Within forest, permeability increased with both forest condition and correspondence with the focal forest type:

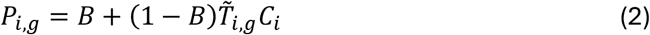

where 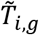 is a transformed measure of forest-type correspondence that reduces differences among forest types, allowing non-focal forest to remain relatively permeable. The transformation and associated numerical implementation are described in Appendix S1. Permeability was highest where forest type matched the focal ecological group and forest condition was high, while non-forest retained the baseline permeability (Fig. 1).

Forest condition therefore influences landscape functionality through two linked pathways: by changing the contribution of a cell as habitat through *Q*_*i,g*_, and by changing the ease of movement through that cell through *P*_*i,g*_. These are not independent perturbations, but two consequences of variation in the same underlying forest-condition variable.

### Movement and ecological distance

Movement between habitat locations was represented using randomized shortest-path (RSP) expected cost distance (Saerens et al. 2009; Van Moorter et al. 2021). Least-cost path models concentrate movement along the lowest-cost route (Adriaensen et al. 2003), whereas random-walk approaches distribute movement across many possible routes (McRae & Beier 2007). The RSP framework spans the continuum between these extremes by allowing movement to depend on both route cost and the availability of alternative pathways (Van Moorter et al. 2021). We used a fixed intermediate parameterization balancing cost and number of alternative pathways (optimality parameter *θ* = 1.0).

For each ecological group *g*, the habitat-permeability surface *P*_*i,g*_ was converted to movement cost (−log (*x*)) and used to calculate RSP expected cost between locations. The resulting accumulated movement costs were rescaled to meter-equivalent units using their relationship with Euclidean distance under baseline matrix permeability (Appendix S2). We denote this ecological distance between locations *s* and *t* as *d*_*st*_.

Ecological distance was converted to pairwise connection strength using an exponential decay kernel,

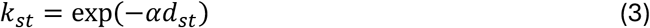

where *k*_*st*_ is the connection strength between locations *s* and *t, d*_*st*_ is ecological distance in meter-equivalent units, and *α* is the exponential decay rate. We evaluated *α* = 1/1000 and 1/2500, corresponding to characteristic movement scales of 1000 and 2500 m-equivalent (Appendix S2). Smaller values of *α* therefore represent broader-scale movement, whereas larger values represent more local movement.

To make the national analysis computationally tractable, ecological distances and pairwise connection strengths were evaluated within a 5-km moving window around blocks of focal cells. This limits connectivity calculations to the surrounding local landscape and therefore truncates the exponential dispersal kernel at the window boundary. For the broader movement scale, connection strength had not declined to negligible values at this distance, such that the resulting estimates represent connectivity within a deliberately restricted spatial neighbourhood rather than the full theoretical extent of the dispersal kernel.

### Landscape functionality

We denote landscape-scale habitat functionality generically as *K*(*L*), where (*L*) represents the landscape (Van Moorter et al. 2023a). In this study, landscape functionality was quantified using the summation-based formulation

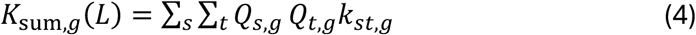

where *Q*_*s,g*_ and *Q*_*t,g*_ are the habitat qualities of locations *s* and *t*, and *k*_*st,g*_ is their pairwise connection strength. The metric therefore integrates habitat amount, habitat quality, and connectivity within a single landscape-scale measure.

Because *K*_*sum,g*_*(L*) scales quadratically with habitat amount under full connectivity, we also expressed landscape functionality as Equivalent Connected Habitat (ECH; Saura et al. 2011; Van Moorter et al. 2023a),

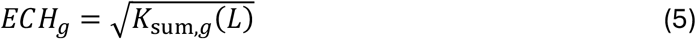

which places the metric on a scale comparable to habitat amount and facilitates interpretation across landscapes.

Landscape functionality was calculated separately for each ecological group. For forest-wide summaries, we additionally defined an equally weighted multi-community landscape functionality as the sum across the spruce-, pine-, and broadleaf-associated networks within each movement scale,

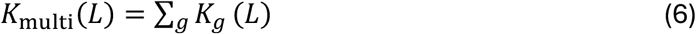

This quantity represents the combined landscape functionality of the three modelled ecological groups and should not be interpreted as a single ecological network.

### Sensitivity of landscape functionality to forest condition

We quantified how a local change in forest condition affects landscape-scale habitat functionality using its analytical sensitivity to *C*_*i*_. For spatial unit *i* and ecological group *g*, sensitivity was defined as

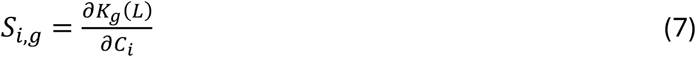

where positive values indicate that increasing forest condition at location *i* increases landscape functionality.

Because forest condition affects both habitat quality and habitat permeability, its effect on landscape functionality can be decomposed into two pathways,

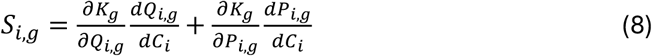

The first term represents the effect of changing the contribution of location *i* as habitat, whereas the second represents the effect of changing its permeability to movement. These components are not independent perturbations, because both arise from a change in the same underlying forest-condition variable. Their sum gives the total sensitivity of landscape functionality to local condition.

For the habitat-quality relationship in Eq. 1,

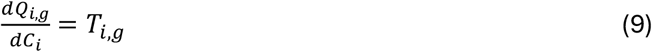

while from Eq. 2,

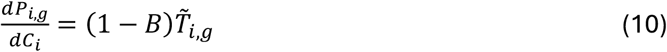

The analytical expressions for ∂*K*/∂*Q* and ∂*K*/∂*P* follow Van Moorter et al. (2026) and are not repeated here. This allows *S*_*i,g*_ to be calculated without explicitly perturbing each cell and recomputing landscape functionality.

Sensitivity was calculated separately for each ecological group and movement scale. It therefore measures the marginal effect of a local change in forest condition in the current landscape configuration, rather than an intrinsic property of the cell itself.

Sensitivity was formulated with respect to landscape functionality *K*(*L*). For the summation-based formulation used here, the corresponding sensitivity of Equivalent Connected Habitat is

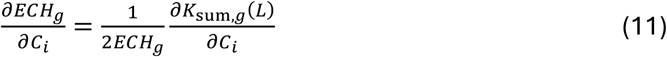

The same marginal change in *K*_sum_ therefore produces a smaller change in ECH where existing ECH is greater. ECH consequently exhibits an intrinsic diminishing-return relationship, whereas sensitivity on the *K* scale remains linear with respect to the underlying change in landscape functionality. We therefore use *K*(*L*) as the primary scale for sensitivity, gain potential, and loss potential, while using ECH as a transformation of landscape functionality into more easily interpretable spatial units.

### Gain and loss potential

Sensitivity quantifies the marginal effect of changing forest condition at a location, but does not account for the current condition of that location. To distinguish the ecological consequences of potential improvement and degradation, we combined sensitivity to obtain first-order estimates of the gains and losses associated with the amount of condition that could respectively be gained or lost.

For spatial unit *i* and ecological group *g*, ‘gain potential’ was defined as

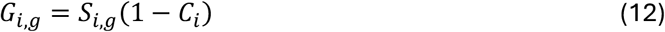

where 1 − *C*_*i*_ represents the current deficit in forest condition, or degradation, corresponding to the maximum possible increase in forest condition relative to the upper bound of the condition scale. Gain potential is therefore high where improving local condition would strongly increase landscape functionality and where substantial scope for improvement remains.

Similarly, ‘loss potential’ was defined as

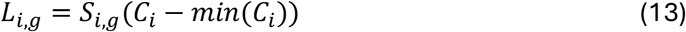

where *C*_*i*_ − *min*(*C*_*i*_) represents the amount of current condition that could potentially be lost (*min*(*C*_*i*_) = 0.2, *see Fig*. 1). Loss potential is therefore high where degradation of an already high-condition location would strongly reduce landscape functionality.

Gain and loss potential were calculated separately for each ecological group and movement scale. For forest-wide summaries and visualizations, we additionally constructed an equally weighted multi-community indicator by summing gain and loss potential across the spruce-, pine-, and broadleaf-associated ecological groups within each movement scale:

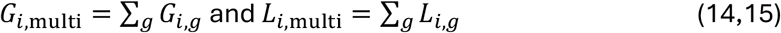

These aggregated values represent the combined contribution of the three modelled ecological communities and should not be interpreted as the gain or loss potential of a single ecological network.

Gain and loss potential represent ecological potential only. They do not incorporate restoration feasibility, management cost, land ownership, threat, or other factors that would be required to translate ecological potential into operational restoration or conservation priorities.

### Protected-area representation

To assess how ecological gain and loss potential are represented within existing protected areas, we overlaid the spatial outputs with the Naturvernområder dataset from Naturbase (Miljødirektoratet 2020). We retained national parks and nature reserves, which provide comparatively strong long-term protection relevant to forest ecosystems, and excluded other protection categories that do not imply comparable restrictions on forest management.

For each ecological group and movement scale, we quantified the distribution of loss and gain potential inside and outside protected areas. We calculated the proportion of total loss and gain potential occurring within protected areas and compared these values with the overall proportion of forest area under protection. This provided a baseline for assessing whether ecological loss and gain potential were disproportionately represented within existing protected areas.

### Software and implementation

Spatial preprocessing, raster calculations, and statistical analyses were conducted in R 4.4.1 (R Core Team 2024) using packages including *terra, sf*, and *ggplot2*. Connectivity and sensitivity calculations were implemented in Julia v1.10 (Bezanson et al. 2017) using *ConScape* (Van Moorter et al. 2023b).

## Results

### Landscape functionality and movement corridors

Forest condition and landscape functionality showed broad but non-identical spatial patterns across Norway (Pearson’s *r=0.30*; Fig. 3). Mean forest condition varied regionally, whereas Equivalent Connected Habitat (ECH) additionally reflected habitat amount, forest-type composition, and connectivity within each 10 × 10 km reporting cell. Areas with the highest forest condition therefore did not necessarily correspond to areas with the highest landscape functionality.

**Figure 3.**
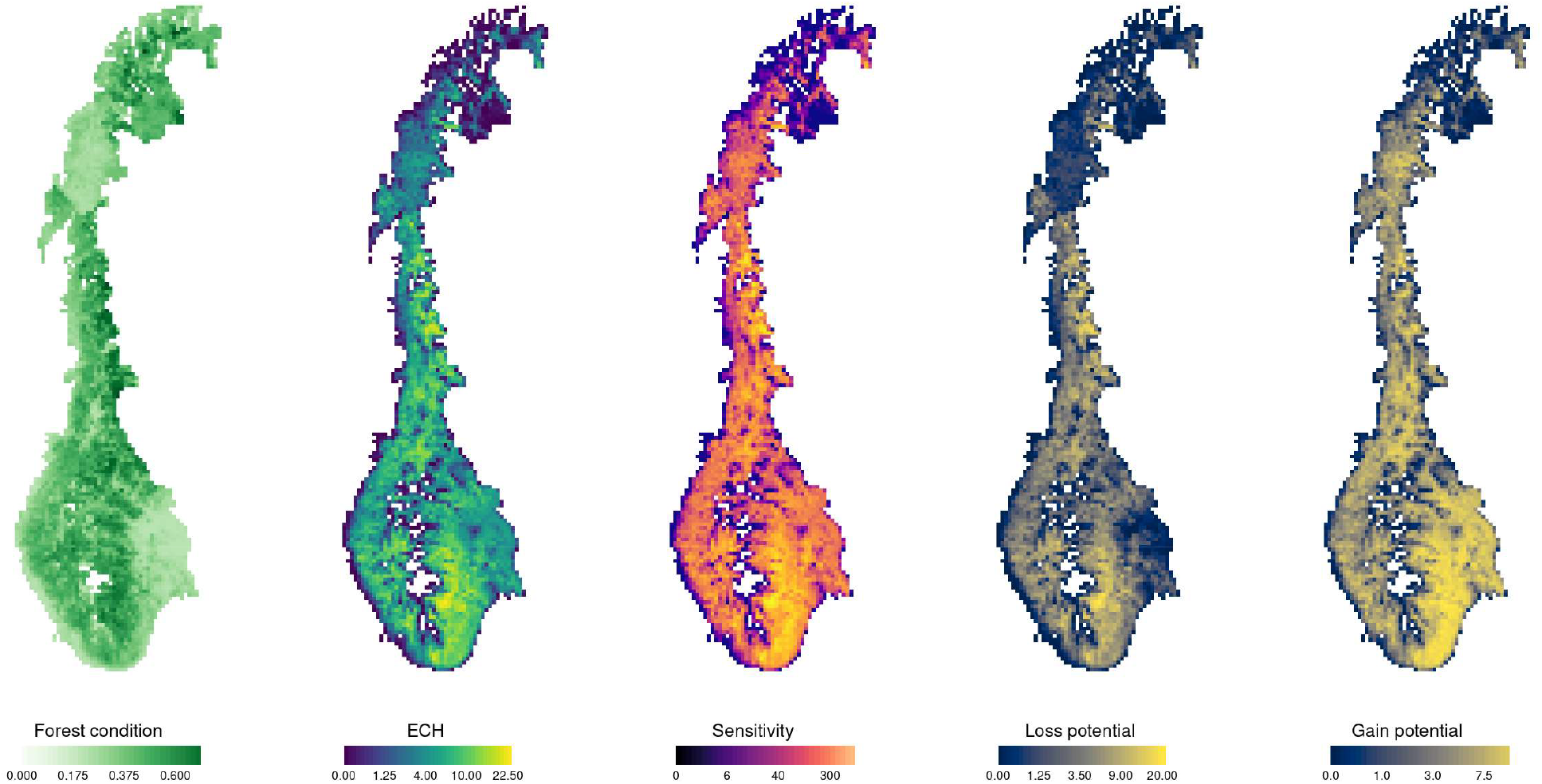
National patterns in forest condition, landscape functionality, sensitivity, and ecological loss and gain potential in Norway. Outputs calculated at 50 × 50 m resolution were summarized within 10 × 10 km grid cells for visualization. Forest condition and sensitivity are shown as means across 50 × 50 m cells within each reporting cell, whereas landscape functionality and loss and gain potential are aggregated over the reporting cell. Landscape functionality is expressed as Equivalent Connected Habitat (ECH; km^2^). Sensitivity is shown as the mean change in landscape functionality K per unit change in forest condition. Loss and gain potential were calculated from sensitivity on the K scale and converted to local changes in ECH using the current landscape ECH within each 10 × 10 km cell as baseline. These ECH-change values represent local potential changes and are not additive across reporting cells. Values are aggregated across the spruce-, pine-, and broadleaf-associated ecological groups for the small-scale movement scenario.

To illustrate the fine-scale spatial structure underlying these national patterns, we selected a focal landscape in southern Norway and visualized results for the spruce-associated ecological group under the local movement-scale scenario (Fig. 4). The focal landscape was used for spatial interpretation only; all summaries and comparisons were based on the complete, high-resolution, national analysis.

**Figure 4.**
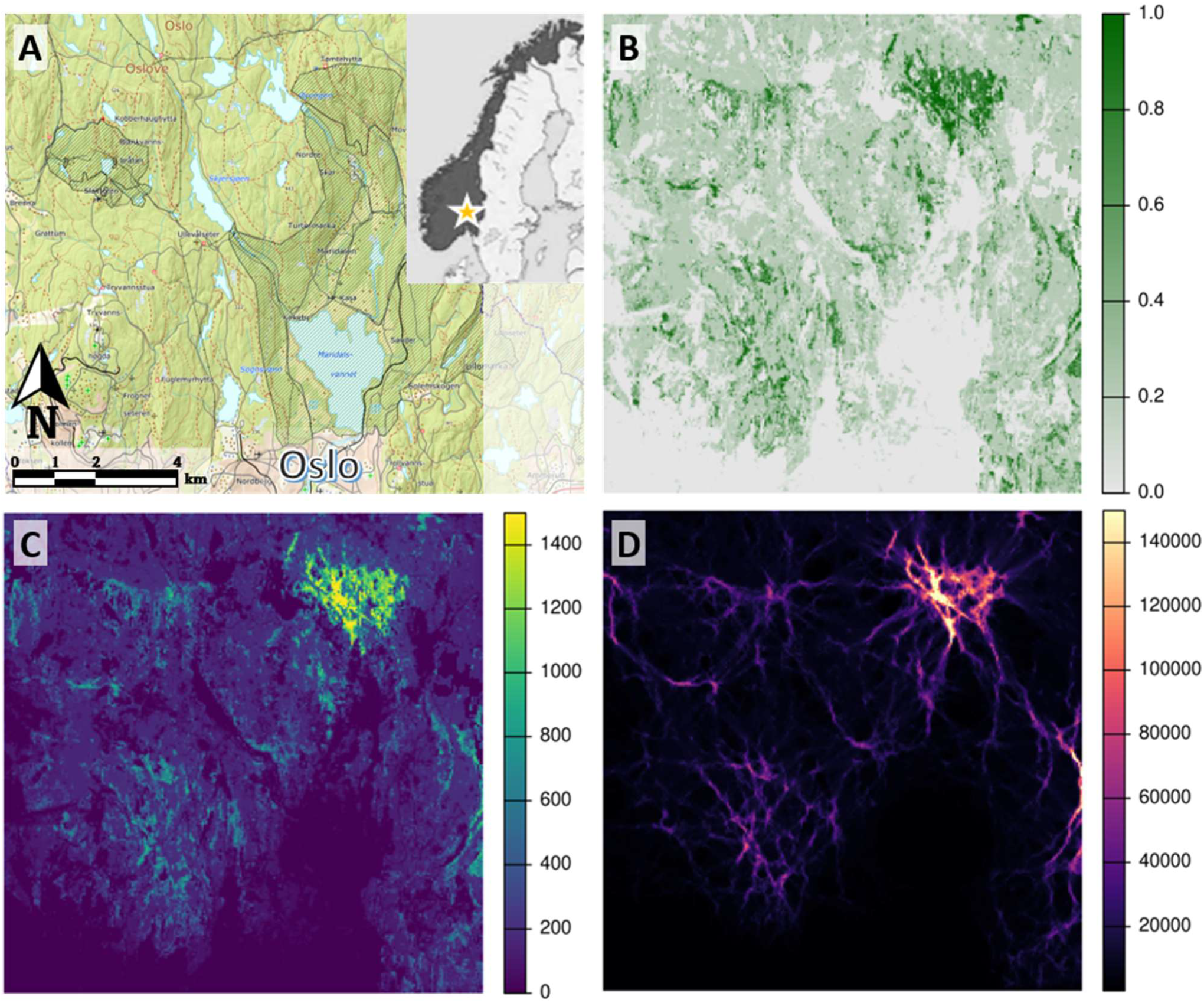
Fine-scale illustration of habitat quality, habitat functionality, and corridors for the spruce-associated ecological group under the local movement-scale scenario. A) Location of the focal landscape in Norway, north of Oslo. B) Spruce habitat quality, calculated from forest condition and the local proportion of spruce, showing the spatial distribution of habitat available to the focal ecological group. C) Habitat functionality, highlighting cells that combine high habitat quality with strong connectivity to surrounding habitat. D) Corridor value, highlighting cells that contribute strongly to movement between habitat areas. The focal landscape is shown for spatial interpretation; national summaries were based on the full national analysis.

Spruce habitat quality, which combines forest condition with the local proportion of spruce, was spatially heterogeneous across the focal landscape (Fig. 4B) with one larger patch of high-condition spruce forest within the ‘Mellomkollen’ nature reserve and several smaller patches scattered across the area. Habitat functionality highlights the high-condition forests that are well-connected to other high-condition forests; it therefore emphasizes the large patch of high-condition forest compared to the smaller fragmented patches (Fig. 4C).

Corridor values showed a complementary spatial pattern. High corridor values frequently occurred within areas of high habitat functionality, reflecting the fact that well-connected habitat can also form part of important movement pathways. However, the two metrics were not equivalent. Some corridor cells occurred outside areas of highest habitat functionality, while substantial areas of highly functional habitat did not form major movement pathways (Fig. 4C–D). Across the national analysis, habitat functionality and corridor value were strongly associated, but not redundant (Spearman’s *r*_*s*_ *= 0.78*), with 62.9% overlap among the top 1% of cells and 72.4% among the top 5%. Thus, corridor value identified a movement-related role that was not identical to the habitat functionality. Both functional habitat and corridors showed highly similar patterns across movement scales (Pearson’s *r=0.SS* and *r=0.S8*, respectively).

### Sensitivity through habitat quality and permeability

Mean sensitivity within each 10 × 10 km reporting cell showed a spatial pattern distinct from forest condition (Pearson’s *r=0.22*; Fig. 3), but was closely associated with current landscape functionality (*r =0.S0*; Fig. 3). This strong relationship was expected because sensitivity is greatest where locations are well connected to high-quality habitat or occupy important corridor positions, and such configurations are most prevalent in well-functioning ecological networks. High-sensitivity locations were those where small changes in forest condition were expected to produce relatively large changes in landscape functionality.

Within the focal landscape, the two pathways through which forest condition affected landscape functionality showed broadly similar spatial patterns, but differed in their local emphasis (Fig. 5). Sensitivity through habitat quality was relatively broad and diffuse, identifying cells where changes in forest condition would directly alter the contribution of spruce habitat to landscape functionality (Fig. 5A). In contrast, sensitivity through habitat permeability was more spatially concentrated and aligned with corridor-like structures and bottlenecks (Fig. 5B), identifying locations where changes in condition would strongly affect movement through the surrounding network. Across the national analysis, the two sensitivity components were positively associated but not redundant (Spearman’s *r*_*s*_ *= 0.C2*); only 28.1% of cells in the top 1% for one mechanism were also in the top 1% for the other, increasing to 39.6% for the top 5%. Van Moorter et al. (2026) show the mathematical relationship between quality-based sensitivity and functional habitat, and between permeability-based sensitivity and corridors, which explains the similarity in their spatial patterns in the focal landscape.

**Figure 5.**
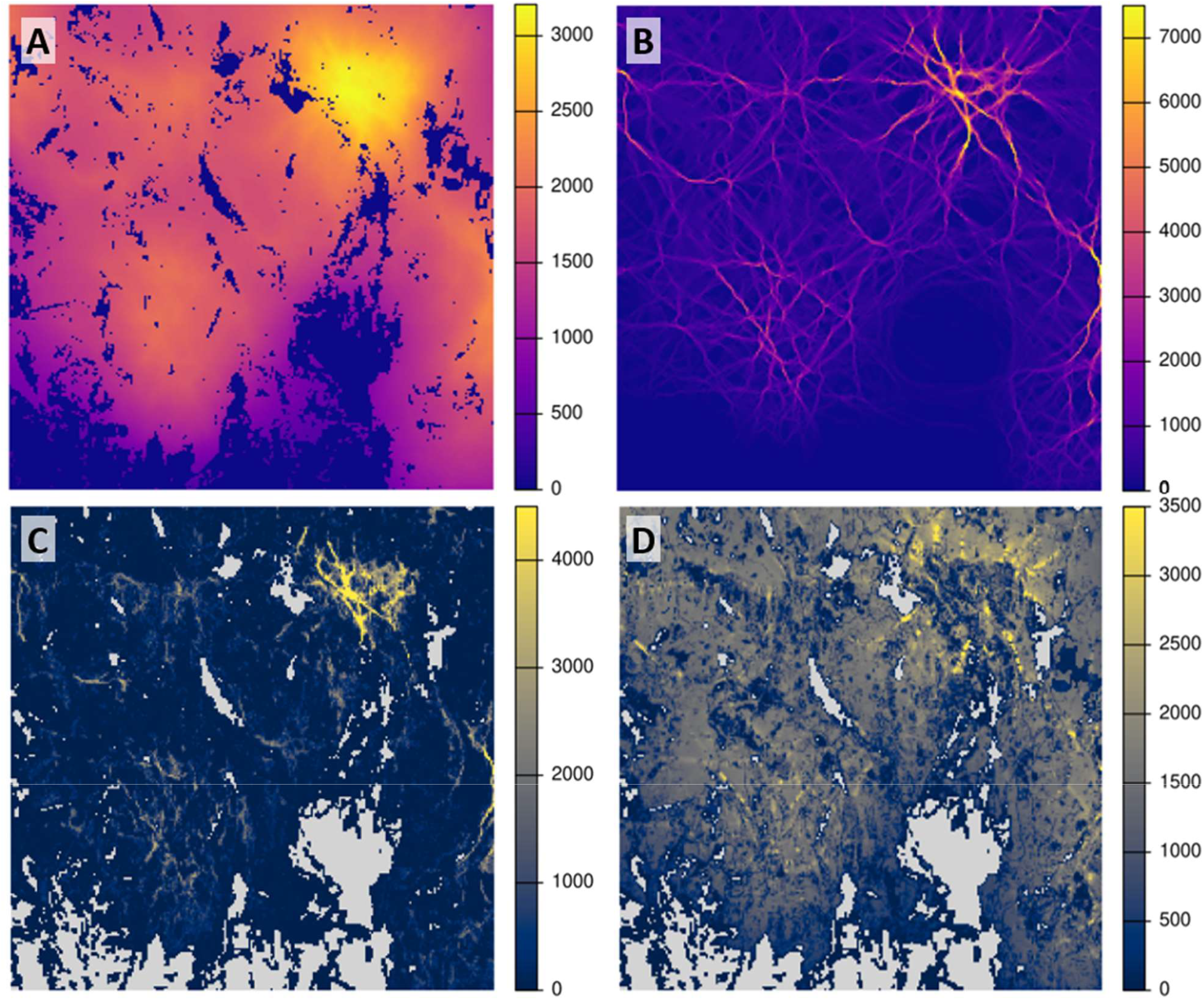
Sensitivity-based loss and gain potential for the same focal landscape shown in Figure 4. A) Sensitivity of landscape functionality K(l) to changes in habitat quality. High values indicate cells where increasing habitat quality would most increase the local habitat contribution to connected natural spruce forest. B) Sensitivity of landscape functionality K(l) to changes in permeability. High values indicate cells where changes in habitat permeability would most affect movement and connectivity through the landscape, often along corridor-like structures. C) Loss potential, calculated as sensitivity weighted by current forest condition, highlighting high-condition areas where loss or degradation would have a large effect on landscape functionality. D) Gain potential, calculated as sensitivity weighted by forest degradation (i.e. condition deficit), highlighting lower-condition areas where improvement in forest condition would be expected to produce large gains in landscape functionality. All outputs were calculated as part of the national analysis and are shown here as a focal-area visualization.

The difference in spatial distribution between the two sensitivity pathways in the focal area was also reflected in how their contributions accumulated across Norway (Fig. 6). Permeability-based sensitivity accumulated more rapidly among the highest-ranked cells, indicating that its largest effects were concentrated in a relatively small fraction of the landscape. As progressively lower-ranked cells were included, additional permeability sensitivity contributed proportionally less, whereas habitat-quality sensitivity continued to accumulate across a broader fraction of the landscape and ultimately reached a larger total contribution. This pattern was evident at both movement scales. Cumulative permeability sensitivity was relatively similar between the local- and broad-movement scenarios, whereas cumulative habitat-quality sensitivity was substantially greater under broad movement. The ranking of cells according to quality and permeability sensitivity was highly similar across movement scales (Spearman’s *r*_*s*_*=0.S8* and *r*_*s*_*=0.S8*, respectively).

**Figure 6.**
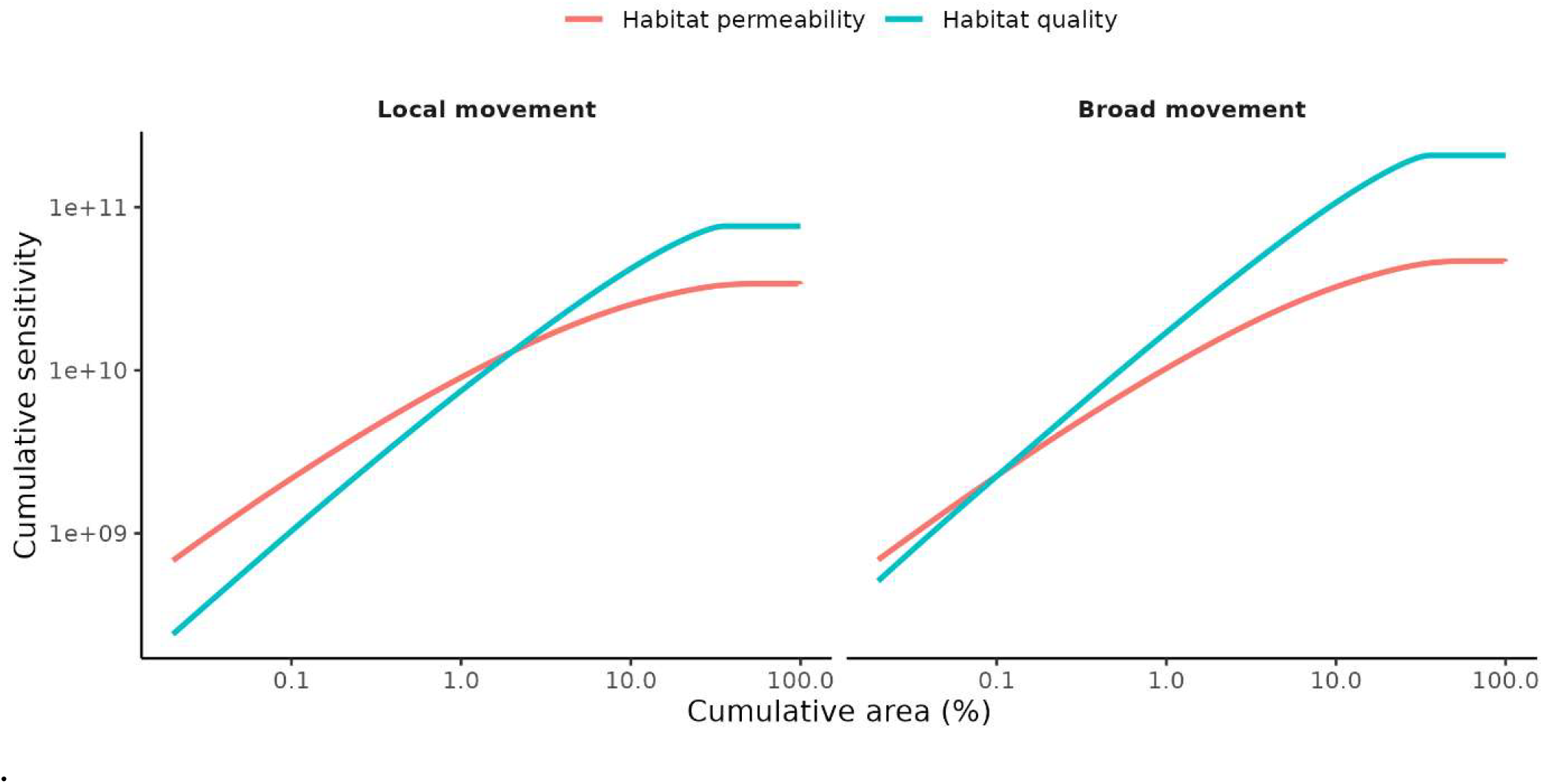
Cumulative contribution of habitat-quality and habitat-permeability sensitivity across progressively larger fractions of the landscape. Cells were ranked separately by their sensitivity through habitat quality and habitat permeability, and cumulative sensitivity was calculated as progressively lower-ranked cells were included. Curves are shown for the local and broad movement-scale scenarios and are aggregated across the spruce-, pine-, and broadleaf-associated ecological groups. Both axes are shown on logarithmic scales. Habitat-permeability sensitivity accumulated more rapidly among the highest-ranked cells, indicating stronger concentration in a small fraction of the landscape, whereas habitat-quality sensitivity accumulated to a larger total contribution as progressively larger areas were included.

### Ecological loss and gain potential and their representation in protected areas

Combining sensitivity with the amount of forest condition that could be lost or gained produced distinct national patterns of ecological loss and gain potential (Fig. 3). Loss potential was highest where relatively high current condition coincided with high sensitivity, whereas gain potential emphasized areas where sensitivity remained high but current forest condition was lower, leaving greater scope for improvement.

Within the focal landscape, the loss and gain potential showed distinctly different patterns (Fig. 5CCD). Also across Norway, the ranking across loss and gain potential differed, albeit positively associated (Spearman’s *r*_*s*_*=0.38*). Areas with high loss potential showed limited overlap with areas of high gain potential, with 7.6% overlap among the top 1% of cells and 19% among the top 5%. Loss potential was strongly positively associated with forest condition (*r*_*s*_ *=0.7S*), and high-loss cells frequently coincided with high-condition forests (18% overlap among the top 1% and 41% among the top 5%). By contrast, gain potential showed little association with forest degradation (*r*_*s*_ *=0.08*), and high-gain cells showed very little overlap with the most degraded forests, with 0.1% among the top 1% and 3% among the top 5%. Thus, areas where restoration would produce the largest gains were not necessarily the same areas where degradation would have the largest ecological consequences, nor were they simply the most degraded forests. The ranking of cells according to loss and gain potential was highly similar across movement scales (Spearman’s *r*_*s*_*=1.00* and *r*_*s*_*=0.S8*, respectively). Figure 5D shows areas with high gain potential around the patch of high-condition spruce forest in the northeast of our focal area. This illustrates that high gain potential can occur in areas that are currently degraded but remain well connected to high-condition forest or lie along corridors between such forests, reflecting high sensitivity through habitat quality or permeability, respectively.

Existing protected areas represented only a small fraction of the highest-ranked ecological loss and gain potential (Fig. 7). Representation was greatest for the very highest-ranked loss-potential cells (top 0.01%), of which approximately 21% occurred within national parks or nature reserves. However, coverage declined rapidly as progressively larger fractions of the landscape were included, falling to around 10% for the top 10% of ranked area. Gain potential was consistently less well represented than loss potential: 13% of the top 1% of loss-potential cells were protected compared with 7% of the top 1% of gain-potential cells, and corresponding values for the top 5% were 11% and 6%. The percentage of loss and gain potential that falls within protected areas shows a similar pattern across forest types and movement scales; loss potential is about twice as well protected as gain potential (Table 2; 7-12% versus 4-5%, respectively). Loss potential for spruce forests is slightly better protected than pine and broadleaf (12, 9, and 8%, respectively); the differences among movement scales were minimal (less than 1%). Thus, although the highest-loss areas received somewhat greater protection than the highest-gain areas, both were generally poorly represented within existing protected areas.

**Table 1.**

| Term | Symbol | Definition / interpretation |
| --- | --- | --- |
| Habitat condition | $N_i$ | Local ecological state of spatial unit $i$; represented here by forest naturalness. |
| Forest type / ecological group | $g$ | Broad forest category used to define separate ecological networks; here spruce-, pine-, and broadleaf-dominated forest. |
| Tree-species proportion | $T_{i,g}$ | Proportion of spatial unit $i$ associated with forest type $g$. |
| Habitat quality | $Q_{i,g}$ | Contribution of spatial unit $i$ as habitat for ecological network $g$, determined by forest type and habitat condition. |
| Landscape permeability | $P_{i,g}$ | Ease of movement through spatial unit $i$ for ecological network $g$. |
| Ecological distance | $d_{st}$ | Effective distance between locations $s$ and $t$ after accounting for permeability of the intervening landscape. |
| Proximity / connection strength | $k_{st}$ | Strength of connection between locations $s$ and $t$, derived from ecological distance. |
| Landscape functionality | $K(L)$ | Generic landscape-level measure integrating habitat amount/quality and connectivity; abbreviated as landscape functionality. Different formulations can be used depending on how landscape structure is summarized. |
| Summation-based landscape functionality | $K_{\text{sum}}(L)$ | Landscape functionality obtained by summing pairwise habitat contributions weighted by connectivity. This is the formulation underlying ECH in this study. |
| Eigenvalue-based landscape functionality | $K_{\text{eig}}(L)$ | Landscape functionality based on the leading eigenvalue of the landscape matrix; corresponds to metapopulation-capacity-type formulations. |
| Equivalent Connected Habitat | $ECH$ | $ECH = \sqrt{K_{\text{sum}}(L)}$; expresses landscape functionality as an equivalent amount of fully connected habitat. |
| Sensitivity | $S_i = \frac{\partial K}{\partial N_i}$ | Response of landscape functionality to a small change in habitat condition at location $i$. |
| Loss potential | $C_i = S_i N_i$ | Potential ecological loss associated with degradation of habitat condition at location $i$. |
| Gain potential | $R_i = S_i(1 - N_i)$ | Potential ecological gain associated with improvement of habitat condition at location $i$. |
| Conservation / restoration priority | — | Broader management priorities that may combine loss or gain potential, respectively, with threat, feasibility, cost, ownership, governance, and other constraints. |
| Ecological network | — | Habitat locations and functional connections representing a focal forest-associated community or model group. |
| Spatial unit | $i, j, s, t$ | Generic local unit of analysis; in this application, a raster cell. |

**Table 2.** Protected-area representation of loss and gain potential by movement scale and forest type. For each potential objective, the table reports the total volume across the national forest landscape, the volume occurring inside protected areas, and the corresponding protected share. Values are shown separately for loss and gain potential, for short- and long-distance movement assumptions, and for spruce, pine, and broadleaf forest networks, with multi-community aggregation across forest groups. Loss potential was more strongly represented within protected areas than gain potential across all movement scales and forest-group networks.

| forest group | scale | total loss volume | protected loss volume | protected loss % | total gain volume | protected gain volume | protected gain % |
| --- | --- | --- | --- | --- | --- | --- | --- |
| spruce | small | 6.97E+09 | 8.87E+08 | 12.73 | 1.11E+10 | 5.54E+08 | 5.00 |
| pine | small | 6.02E+09 | 5.90E+08 | 9.79 | 1.15E+10 | 5.92E+08 | 5.14 |
| broadleaf | small | 9.21E+09 | 7.11E+08 | 7.71 | 1.02E+10 | 4.25E+08 | 4.18 |
| community | small | 2.22E+10 | 2.19E+09 | 9.85 | 3.27E+10 | 1.57E+09 | 4.80 |
| spruce | large | 1.32E+10 | 1.58E+09 | 11.96 | 2.82E+10 | 1.19E+09 | 4.23 |
| pine | large | 1.28E+10 | 1.17E+09 | 9.11 | 2.88E+10 | 1.31E+09 | 4.54 |
| broadleaf | large | 1.63E+10 | 1.10E+09 | 6.74 | 2.34E+10 | 8.61E+08 | 3.68 |
| community | large | 4.23E+10 | 3.85E+09 | 9.09 | 8.04E+10 | 3.36E+09 | 4.18 |

**Figure 7.**
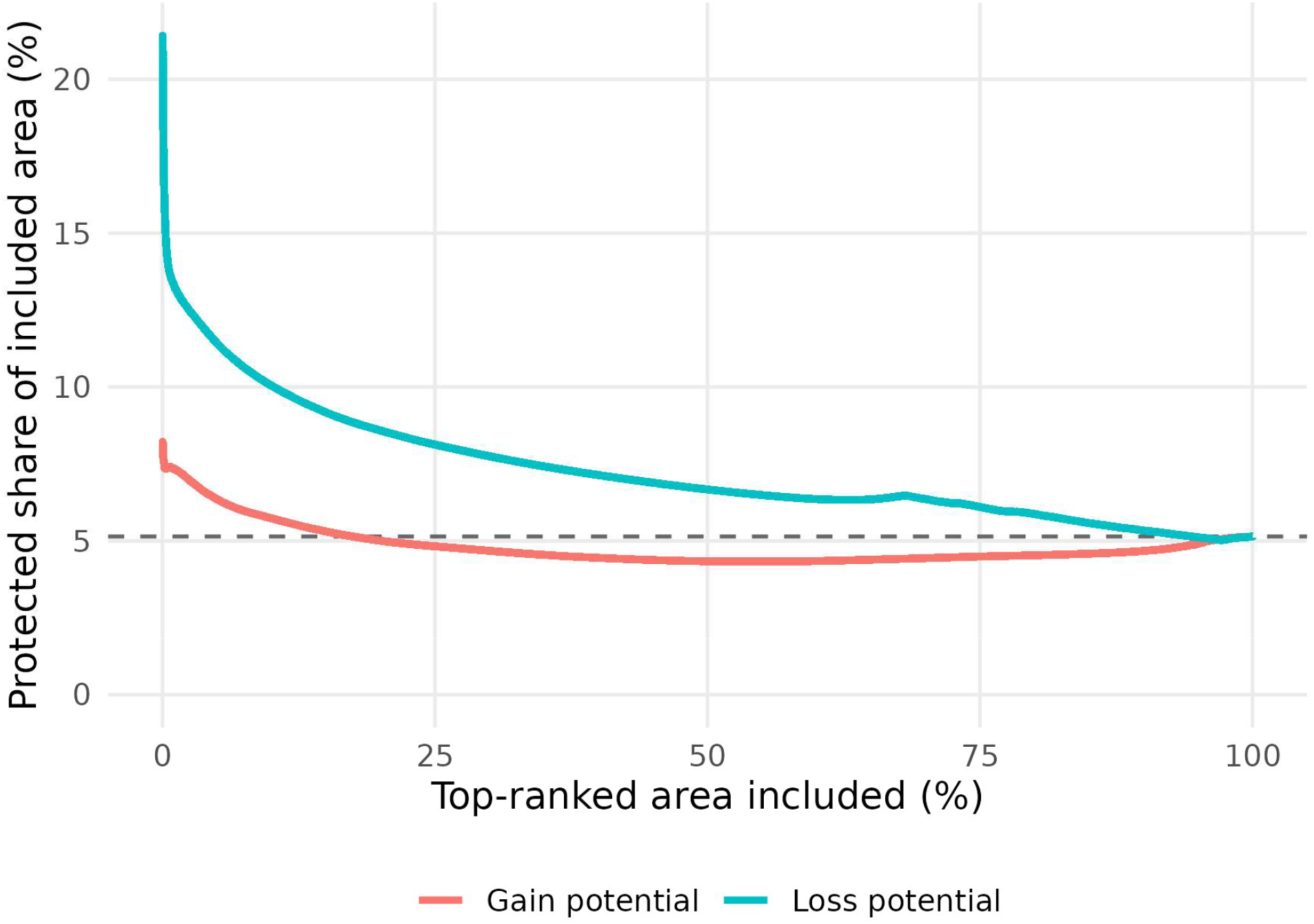
Protected-area coverage of top-ranked areas with highest loss and gain potential. Curves show the proportion of top-ranked loss-potential and gain-potential areas that fall within protected areas as progressively larger fractions of the landscape are included. Areas were ranked separately from highest to lowest loss or gain potential. The dashed horizontal line indicates the overall protected share of the Norwegian forest area. Loss potential is more strongly represented within protected areas than expected from the landscape-wide protection level, especially among the highest-ranked areas. Gain potential is less well represented than loss potential across most of the ranking distribution, although the highest-ranked gain-potential areas are also represented above the landscape-wide baseline. Overall, both loss and gain potential are only partly captured by the current protected-area network, with gain potential generally less well represented than loss potential.

## Discussion

We extended perturbation-based approaches for spatial prioritization by using analytical sensitivity to identify where changes in habitat condition are expected to have the greatest effect on persistence-relevant landscape functionality. Following sensitivity-based conservation prioritization (Van Moorter et al. 2026), we treat both degradation and restoration as perturbation problems: where would a local change in habitat condition most strongly affect connected habitat? Applied to Norwegian forest landscapes, this framework quantified ecological loss and gain potential from a common underlying sensitivity surface while distinguishing effects mediated through habitat quality and landscape permeability. The results show that forest condition, sensitivity, and ecological loss and gain potential are related but not interchangeable. In particular, areas with the greatest gain potential were neither simply the most degraded forests nor the same areas where degradation would cause the greatest losses. These results show that ecological leverage and current condition represent distinct dimensions of prioritization: sensitivity identifies where change matters most, while current condition determines whether that leverage translates into loss or gain potential.

The ecological importance of a location depends not only on its local condition but also on its position within the surrounding habitat network, because local habitat characteristics and landscape context jointly influence dispersal, colonization, and population persistence (Frey et al. 2012; Gilby et al. 2018). Connectivity-based approaches for spatial prioritization have therefore increasingly incorporated habitat amount, proximity, network structure, or corridors (Daigle et al. 2020; Tambosi et al. 2014; Ortega et al. 2023; Bubnicki et al. 2024; Wang et al. 2025). Perturbation-based approaches build on this by asking how local changes in habitat condition alter persistence-relevant measures of connected habitat (Ovaskainen 2003; Ovaskainen & Hanski 2003; Saura & Torné 2009; García-Feced et al. 2011; Foltête et al. 2014), such as metapopulation capacity (Hanski & Ovaskainen 2000) or probability of connectivity (Saura & Pascual-Hortal 2007). Repeatedly recalculating such metrics for local perturbations becomes computationally demanding in large, high-resolution landscapes. Analytical sensitivity provides a scalable alternative by estimating the marginal effect of a local change directly from the landscape model (Van Moorter et al. 2026). Importantly, sensitivity itself is direction-neutral: it identifies where landscape functionality is most responsive to changes in condition. Whether this ecological leverage translates into loss or gain potential depends on current habitat condition. High sensitivity combined with high current condition produces high loss potential, whereas high sensitivity combined with substantial scope for ecological improvement produces high gain potential.

The framework is applicable across ecosystems for which habitat condition can be linked to network properties, and we demonstrate this general approach in Norwegian forest landscapes. Separating habitat-quality- and permeability-mediated sensitivity provides an ecologically interpretable decomposition of how local condition affects landscape-scale connected habitat. Habitat-quality sensitivity was relatively broad and diffuse, identifying locations where improved condition would directly increase the amount and quality of habitat contributing to landscape functionality. Permeability sensitivity was more spatially concentrated and associated with corridor-like structures and bottlenecks, where relatively small changes could influence movement among habitat areas. This distinction is consistent with the broader recognition that habitat areas and the connections among them can make different contributions to landscape connectivity and habitat availability (Saura & Rubio 2010; Baguette et al. 2013). Although habitat functionality and corridor value were strongly associated in our analysis, and the two sensitivity pathways were positively correlated, their limited overlap among the highest-ranked locations shows that habitat provision and movement support are not interchangeable landscape functions. The concentrated distribution of permeability sensitivity also meant that a relatively small fraction of the landscape accounted for a large proportion of its total contribution, whereas habitat-quality sensitivity was distributed more broadly. The relative importance of permeability was greater under the more localized movement scenario, consistent with the expectation that the intervening landscape becomes increasingly important as dispersal capacity declines (Baguette et al. 2013; Lachat et al. 2025). Improving permeability may therefore generate substantial gains through strategically located interventions, whereas gains through habitat-quality improvement may be distributed across a broader area.

Combining the two sensitivity pathways with current forest condition translated these patterns into ecological loss and gain potential. High loss potential was concentrated in relatively high-condition forest that already contributed strongly to landscape functionality, whereas high gain potential tended to occur where lower-condition forest remained embedded within functional habitat networks, including areas adjacent to high-condition habitat or situated along important movement corridors. Consequently, degradation itself was a poor surrogate for gain potential: gain potential showed little association with forest degradation, and the highest-gain locations showed almost no overlap with the most degraded forest. This result is consistent with restoration frameworks emphasizing that ecological condition needs to be considered together with landscape context, habitat amount, and connectivity when identifying restoration opportunities (Tambosi et al. 2014; Gilby et al. 2018; Brancalion et al. 2019; Strassburg et al. 2020; Wang et al. 2025). Our framework extends this logic by estimating directly how a marginal improvement in condition at each location would alter landscape functionality, rather than treating connectivity or neighbourhood condition solely as static attributes of candidate restoration sites.

The protected-area analysis highlights generally low coverage of areas with high ecological loss and gain potential, but also a clear asymmetry between them. Existing national parks and nature reserves represented high loss potential more strongly than high gain potential. Nevertheless, coverage declined rapidly beyond the very highest-ranked loss-potential locations, and both loss and gain potential were generally poorly represented within current protected areas. The particularly low representation of gain potential suggests that ecological restoration outside protected areas could play an important complementary role by enlarging, buffering, or reconnecting areas of high-condition forest and by improving the permeability of the surrounding matrix. Such an approach is consistent with broader calls to integrate connectivity and restoration into spatial conservation planning beyond reserve boundaries (Daigle et al. 2020; Ortega et al. 2023; Hua et al. 2024).

Loss and gain potential should not, however, be interpreted directly as conservation and restoration priorities. They represent the ecological benefit component of a broader prioritization problem: where preventing a decline or achieving an improvement in habitat condition would have the greatest marginal effect on landscape functionality. Operational priorities additionally depend on whether the relevant change in condition is feasible, how much improvement can realistically be achieved, how long recovery is expected to take, what the intervention costs relative to alternatives, and how it interacts with other ecosystem services and management objectives (Orsi & Geneletti 2010; Suding 2011; Strassburg et al. 2020). These constraints are particularly important in forests because ecological recovery can be slow and may depend on structural attributes and habitat continuity that develop over decades or centuries (Fritz et al. 2008; Majdanová et al. 2023; Hoffmann et al. 2025; Macdonald et al. 2026). A recently clear-cut stand, for example, may have a large theoretical condition deficit, but much of the structure associated with near-natural or old forest may not be recoverable within management-relevant time frames. A less degraded forest with somewhat lower ecological gain potential could therefore provide a more achievable restoration opportunity. Conceptually, the gain potential estimated here can be viewed as an ecological opportunity layer that could subsequently be combined with an expected restoration-response function describing how much of the condition deficit is realistically recoverable under a particular intervention, together with information on cost, land tenure, risk, and competing objectives. Restoration also need not require active manipulation: where natural recovery processes remain effective, preventing further degradation may allow passive recovery, whereas elsewhere active measures such as increasing dead wood, restoring tree-species composition, or promoting structural heterogeneity may accelerate recovery (Hoffmann et al. 2025). The distinction between ecological potential and achievable restoration benefit is therefore essential when translating sensitivity maps into management decisions.

The choice between *K* and ECH affects how sensitivity and gain and loss potential are interpreted. We used Equivalent Connected Habitat (ECH) because it transforms the underlying summation-based landscape functionality *K* to a scale comparable with habitat amount and is therefore useful for describing the current state of a landscape. Sensitivity, loss potential, and gain potential were evaluated on the underlying *K* scale, except in Fig. 3, where loss and gain potential are reported as local changes in ECH. This distinction matters because ECH is proportional to the square root of *K*. Consequently, an identical increase in *K* translates into a smaller increase in ECH where existing ECH is already high. ECH therefore has an intrinsic diminishing-return relationship, making marginal changes dependent on the baseline landscape state and reporting extent. By contrast, *K* is the direct scale of the underlying landscape-functionality measure and avoids the additional diminishing-return transformation introduced by ECH. We therefore regard *K* as the more natural scale for ranking marginal ecological effects, while ECH remains useful for communicating the amount of connected habitat represented by a specific landscape.

As with other network- and perturbation-based approaches, the results remain conditional on assumptions in the underlying landscape model. These include the representation of habitat quality and permeability, the selected movement kernel and spatial scale, the spatial resolution of the analysis, and the extent to which landscape functionality captures processes relevant to population persistence. Species differ substantially in habitat requirements, dispersal behaviour, and the spatial scales at which landscape structure affects their populations (Baguette et al. 2013; Hunter-Ayad & Hassall 2020; Lachat et al. 2025; Lapin et al. 2025). Our community-level parameterizations therefore represent broad ecological scenarios rather than predictions for individual species. Despite this uncertainty, rankings of sensitivity and loss and gain potential were highly similar across the two movement scales considered here, suggesting that the main spatial patterns were robust within the range of movement assumptions evaluated. The analysis is also dependent on the quality of the underlying spatial data. Errors in maps of forest composition and condition propagate into habitat quality, permeability, sensitivity, and ultimately loss and gain potential. Improving these underlying spatial representations and validating the resulting ecological networks against independent biodiversity data remain important priorities.

Finally, sensitivity describes the derivative of landscape functionality around the current landscape configuration and therefore estimates the effect of a marginal local change rather than the exact consequence of a large intervention. As the magnitude of habitat change increases, nonlinear responses may cause realized changes to diverge from the first-order approximation. Nevertheless, Van Moorter et al. (2026) showed that sensitivity-based loss potential closely reproduced priorities derived from explicit node destruction, indicating that the analytical approximation preserves much of the information relevant for ranking local interventions. Furthermore, as in most perturbation-based approaches (Ovaskainen 2003; Ovaskainen & Hanski 2003; Saura & Torné 2009; García-Feced et al. 2011; Foltête et al. 2014), effects are evaluated for individual perturbations in isolation, whereas interactions among multiple interventions can introduce important non-linearities. When multiple interventions are implemented, their effects may be synergistic or redundant, such that the combined outcome differs from the sum of independently estimated effects (Hamonic et al. 2023). Sensitivity-based rankings are therefore most naturally interpreted as rankings of marginal ecological effects in the current landscape, whereas optimization approaches that explicitly account for cumulative effects may be preferable when selecting a fixed portfolio of interacting interventions (Hamonic et al. 2025).

The sensitivity-based approach retains the ecological logic of perturbation-based prioritization while avoiding the need to repeatedly recalculate the full connectivity metric for every candidate location. By deriving loss and gain potential from the same sensitivity framework, it provides a consistent way to distinguish where degradation would cause disproportionate losses from where ecological improvement could generate disproportionate gains. These ecological potentials address different management questions and should be interpreted as complementary components of broader conservation and restoration prioritization.

## Conclusion

By differentiating a persistence-relevant connectivity metric with respect to habitat condition, sensitivity-based prioritization shifts spatial planning from mapping where ecosystems are degraded or valuable to estimating where local change is expected to matter most. The Norwegian application shows that this approach can be implemented at national scale and can produce interpretable ecological loss and gain layers for both conservation and restoration. More broadly, the framework offers a scalable way to link local habitat-condition change to biodiversity persistence goals and to identify where limited conservation and restoration resources may have the greatest ecological effect.

## References

Adriaensen, F., Chardon, J. P., De Blust, G., Swinnen, E., Villalba, S., Gulinck, H., & Matthysen, E. (2003). The application of ‘least-cost’modelling as a functional landscape model. Landscape and urban planning, 64(4), 233–247.

Artsdatabanken. (2021). Status for truete arter i skog. Norsk rødliste for arter 2021. Artsdatabanken, Trondheim, Norway.

Asplund, J., Nordén, J., Kjønaas, O. J., Madsen, R. L., Lunde, L. F., Birkemoe, T., … & Nybakken, L. (2024). Long term effects of forest management on forest structure and dead wood in mature boreal forests. Forest Ecology and Management, 572, 122315.

Astrup, R., Rahlf, J., Bjørkelo, K., Debella-Gilo, M., Gjertsen, A. K., & Breidenbach, J. (2019). Forest information at multiple scales: development, evaluation and application of the Norwegian forest resources map SR16. Scandinavian Journal of Forest Research, 34(6), 484–496. 10.1080/02827581.2019.1588989

Baguette, M., Blanchet, S., Legrand, D., Stevens, V., & Turlure, C. (2013). Individual dispersal, landscape connectivity and ecological networks. Biological Reviews, 88. 10.1111/brv.12000

Banks-Leite, C., Ewers, R., Folkard-Tapp, H., & Fraser, A. (2020). Countering the effects of habitat loss, fragmentation, and degradation through habitat restoration. 672–676. 10.1016/j.oneear.2020.11.016

Beger, M., Metaxas, A., Balbar, A. C., McGowan, J. A., Daigle, R., Kuempel, C. D., Treml, E. A., & Possingham, H. P. (2022). Demystifying ecological connectivity for actionable spatial conservation planning. Trends in Ecology & Evolution, 37(12), 1079–1091. 10.1016/j.tree.2022.09.002

Betts, M. G., Yang, Z., Hadley, A. S., Smith, A. C., Rousseau, J. S., Northrup, J. M., … & Gerber, B. D. (2022). Forest degradation drives widespread avian habitat and population declines. Nature Ecology & Evolution, 6(6), 709–719.

Bezanson, J., Edelman, A., Karpinski, S., & Shah, V. B. (2017). Julia: A fresh approach to numerical computing. SIAM Review, 59(1), 65–98. 10.1137/141000671

Brancalion, P. H., Niamir, A., Broadbent, E., Crouzeilles, R., Barros, F. S., Almeyda Zambrano, A. M., … & Chazdon, R. L. (2019). Global restoration opportunities in tropical rainforest landscapes. Science advances, 5(7), eaav3223.

Breidenbach, J., Waser, L.T., Debella-Gilo, M., Schumacher, J., Rahlf, J., Hauglin, M., Puliti, S., Astrup, R., 2021. National mapping and estimation of forest area by dominant tree species using Sentinel-2 data. Can. J. For. Res. 51, 365–379.

Bubnicki, J. W., Angelstam, P., Mikusiński, G., Svensson, J., & Jonsson, B. G. (2024). The conservation value of forests can be predicted at the scale of 1 hectare. Communications Earth & Environment, 5(1), 196.

Burner, R. C., Birkemoe, T., Stephan, J. G., Drag, L., Müller, J., Ovaskainen, O., Potterf, M., Skarpaas, O., Snäll, T., & Sverdrup-Thygeson, A. (2021). Choosy beetles: How host trees and southern boreal forest naturalness may determine dead wood beetle communities. Forest Ecology and Management, 487, 119023. 10.1016/j.foreco.2021.119023

Convention on Biological Diversity. (2022). Kunming-Montreal Global Biodiversity Framework (Decision 15/4, CBD/COP/DEC/15/4).

Daigle, R. M., Metaxas, A., Balbar, A. C., McGowan, J., Treml, E. A., Kuempel, C. D., … & Beger, M. (2020). Operationalizing ecological connectivity in spatial conservation planning with Marxan Connect. Methods in Ecology and Evolution, 11(4), 570–579.

Díaz, S., Zafra-Calvo, N., Purvis, A., Verburg, P. H., Obura, D., Leadley, P., Chaplin-Kramer, R. E., De Meester, L., Dulloo, E., Martín-López, B., Shaw, M. R., Visconti, P., Broadgate, W., Bruford, M. W., Burgess, N. D., Cavender-Bares, J. M., DeClerck, F., Fernández-Palacios, J. M., Garibaldi, L. A., … Zanne, A. E. (2020). Set ambitious goals for biodiversity and sustainability. Science, 370(6515), 411–413. 10.1126/science.abe1530

Edman, M., Gustafsson, M., Stenlid, J., Jonsson, B. G., & Ericson, L. (2004). Spore deposition of wood-decaying fungi: importance of landscape composition. Ecography, 27(1), 103–111.

Esseen, P.-A., Ehnström, B., Ericson, L., & Sjöberg, K. (1997). Boreal forests. Ecological Bulletins, 46, 16–47.

European Parliament, Council of the European Union (2024) Regulation (EU) 2024/1991 of the European Parliament and of the Council of 24 June 2024 on nature restoration and amending Regulation (EU) 2022/869 (Text with EEA relevance). Off J Eur Union, Brussels (BE). http://data.europa.eu/eli/reg/2024/1991/oj

Fahrig, L. (2003). Effects of habitat fragmentation on biodiversity. Annual Review of Ecology, Evolution, and Systematics, 34, 487–515. 10.1146/annurev.ecolsys.34.011802.132419

Foltête, J. C., Girardet, X., & Clauzel, C. (2014). A methodological framework for the use of landscape graphs in land-use planning. Landscape and Urban Planning, 124, 140–150.

Frey, S. J., Strong, A. M., & McFarland, K. P. (2012). The relative contribution of local habitat and landscape context to metapopulation processes: a dynamic occupancy modeling approach. Ecography, 35(7), 581–589.

Fritz, Ö., Gustafsson, L. & Larsson, K. (2008) Does forest continuity matter in conservation? – A study of epiphytic lichens and bryophytes in beech forests of southern Sweden. Biological Conservation, 141, 655–668.

García-Feced, C., Saura, S., & Elena-Rosselló, R. (2011). Improving landscape connectivity in forest districts: A two-stage process for prioritizing agricultural patches for reforestation. Forest ecology and management, 261(1), 154–161.

Gilby, B., Olds, A., Connolly, R., Henderson, C. J., & Schlacher, T. (2018). Spatial Restoration Ecology: Placing Restoration in a Landscape Context. BioScience, 68, 1007–1019. 10.1093/biosci/biy126

Haddad, N. M., Brudvig, L. A., Clobert, J., Davies, K. F., Gonzalez, A., Holt, R. D., Lovejoy, T. E., Sexton, J. O., Austin, M. P., Collins, C. D., Cook, W. M., Damschen, E. I., Ewers, R. M., Foster, B. L., Jenkins, C. N., King, A. J., Laurance, W. F., Levey, D. J., Margules, C. R., … Townshend, J. R. (2015). Habitat fragmentation and its lasting impact on Earth’s ecosystems. Science Advances, 1(2), e1500052. 10.1126/sciadv.1500052

Hägglund, R., & Hjältén, J. (2018). Substrate specific restoration promotes saproxylic beetle diversity in boreal forest set-asides. Forest Ecology and Management, 425, 45–58.

Hamonic, F., Couëtoux, B., Vaxès, Y., & Albert, C. H. (2023). Cumulative effects on habitat networks: How greedy should we be?. Biological Conservation, 282, 110066.

Hamonic, F., Vaxès, Y., Couëtoux, B., & Albert, C. H. (2025). GECOT: Graph-based ecological connectivity optimization tool. Methods in Ecology and Evolution, 16(9), 1914–1922.

Hanski, I. (1999). Metapopulation ecology. Oxford University Press.

Hanski, I., & Ovaskainen, O. (2000). The metapopulation capacity of a fragmented landscape. Nature, 404(6779), 755–758.

Hanson, J. O., Vincent, J., Schuster, R., Fahrig, L., Brennan, A., Martin, A. E., Hughes, J. S., Pither, R., & Bennett, J. R. (2022). A comparison of approaches for including connectivity in systematic conservation planning. Journal of Applied Ecology, 59(10), 2507–2519. 10.1111/1365-2664.14251

Hauglin, M., Asplund, U. J., Jutras-Perreault, M-C., Skarpaas, O., Strimbul, V., Storaunet, K. O., Sverdrup-Thygeson, A., Wold, J. & Ørka, H. O. (2025). Naturskogkart – første versjon. Bak-grunn og metodikk (NIBIO RAPPORT, 11/129). Norsk institutt for bioøkonomi.

Hoffmann, J. A., Teketay, D., Negash, M., & Tesfay, H. M. (2025). Restoring Forest Landscape Connectivity: Why, Where, and How?. In ecological connectivity of forest ecosystems (pp. 265–295). Cham: Springer Nature Switzerland.

Hua, F., Liu, M., & Wang, Z. (2024). Integrating forest restoration into land-use planning at large spatial scales. Current Biology, 34(9), R452–R472.

Hunter-Ayad, J., & Hassall, C. (2020). An empirical, cross-taxon evaluation of landscape-scale connectivity. Biodiversity and Conservation, 29, 1339–1359. 10.1007/s10531-020-01938-2

IPBES. (2019). Global assessment report on biodiversity and ecosystem services of the Intergovernmental Science-Policy Platform on Biodiversity and Ecosystem Services (E. S. Brondízio, J. Settele, S. Díaz, & H. T. Ngo, Eds.). IPBES Secretariat. 10.5281/zenodo.3831673

Jacobsen, R. M., Burner, R. C., Olsen, S. L., Skarpaas, O., & Sverdrup-Thygeson, A. (2020). Near-natural forests harbor richer saproxylic beetle communities than those in intensively managed forests. Forest Ecology and Management, 466, 118124.

Jonsell, M., Hansson, J., & Wedmo, L. (2007). Diversity of saproxylic beetle species in logging residues in Sweden—Comparisons between tree species and diameters. Biological Conservation, 138(1–2), 89–99. 10.1016/j.biocon.2007.04.003

Jonsson, B. G., Kruys, N., & Ranius, T. (2005). Ecology of species living on dead wood—Lessons for dead wood management. Silva Fennica, 39(2), 289–309. 10.14214/sf.390

Kivimäki, I., Van Moorter, B., & Saerens, M. (2024). Sensitivity to network perturbations in the randomized shortest paths framework: theory and applications in ecological connectivity. Journal of Physics: Complexity, 5(2), 025017.

Komonen, A., & Müller, J. (2018). Dispersal ecology of deadwood organisms and connectivity conservation. Conservation Biology, 32(3), 535–545. 10.1111/cobi.13087

Lachat, T., Oettel, J., & Meyer, F. (2025). Do saproxylic species need habitats, connectivity, or connected habitats? In K. Lapin, J. Oettel, M. Braun, & H. Konrad (Eds.), Ecological connectivity of forest ecosystems (pp. 39–54). Springer.

Lapin, K., Konrad, H., Leeb, C., & Oettel, J. (2025). Species on the move: Migration, range shifts, and dispersal of species. In K. Lapin, J. Oettel, M. Braun, & H. Konrad (Eds.), Ecological connectivity of forest ecosystems (pp. 23–38). Springer.

Li, W., Liu, P., Yang, N., Chen, S., Guo, X., Wang, B., & Zhang, L. (2023). Improving landscape connectivity through habitat restoration: application for Asian elephant conservation in Xishuangbanna Prefecture, China.. Integrative zoology. 10.1111/1749-4877.12713

Mace, G. M., Barrett, M., Burgess, N. D., Cornell, S. E., Freeman, R., Grooten, M., & Purvis, A. (2018). Aiming higher to bend the curve of biodiversity loss. Nature Sustainability, 1(9), 448–451. 10.1038/s41893-018-0130-0

Majdanová, L., Hofmeister, J., Pouska, V., Mikoláš, M., Zíbarová, L., Vítková, L., … & Čada, V. (2023). Old-growth forests with long continuity are essential for preserving rare wood-inhabiting fungi. Forest Ecology and Management, 541, 121055.

Margules, C. R., & Pressey, R. L. (2000). Systematic conservation planning. Nature, 405(6783), 243–253.

Matricardi, E. A. T., Skole, D. L., Costa, O. B., Pedlowski, M. A., Samek, J. H., & Miguel, E. P. (2020). Long-term forest degradation surpasses deforestation in the Brazilian Amazon. Science, 369(6509), 1378–1382.

Macdonald, S. E., McIntosh, A. C., Schut, S., Bartels, S., Lee, S. I., Baisheva, E., … & Wu, L. (2026). Biodiversity recovery is slow following clear-cut harvest of boreal forests. Nature Sustainability, 1–11.

McRae, B. H., & Beier, P. (2007). Circuit theory predicts gene flow in plant and animal populations. Proceedings of the National Academy of Sciences, 104(50), 19885–19890.

McRae, B. H., Hall, S. A., Beier, P., & Theobald, D. M. (2012). Where to restore ecological connectivity? Detecting barriers and quantifying restoration benefits. PloS one, 7(12), e52604.

Miljødirektoratet. 2020. Naturvernområder – v2. Norwegian Environment Agency, Norway. Version 30 November 2020. Geonorge Register. https://register.geonorge.no/produktark/naturvernomrader-v2

Moilanen, A., Wilson, K., & Possingham, H. (2009). Spatial conservation prioritization: quantitative methods and computational tools. Oxford University Press.

Noss, R. F., Nielsen, S. E., & Vance-Borland, K. (2009). Prioritizing ecosystems, species, and sites for restoration. In A. Moilanen, K. A. Wilson, & H. P. Possingham (Eds.), Spatial conservation prioritization: Quantitative methods and computational tools (pp. 158–171). Oxford University Press. doi:10.1093/oso/9780199547760.003.0012.

Nordén, J., Penttilä, R., Siitonen, J., Tomppo, E., & Ovaskainen, O. (2013). Specialist species of wood-inhabiting fungi struggle while generalists thrive in fragmented boreal forests. Journal of Ecology, 101(3), 701–712.

Norros, V., Penttilä, R., Suominen, M., & Ovaskainen, O. (2012). Dispersal may limit the occurrence of specialist wood decay fungi already at small spatial scales. Oikos, 121, 961–974. 10.1111/j.1600-0706.2012.20052.x

Östlund, L., Zackrisson, O., & Axelsson, A.-L. (1997). The history and transformation of a Scandinavian boreal forest landscape since the 19th century. Canadian Journal of Forest Research, 27(8), 1198–1206. 10.1139/x97-070

Orsi, F., & Geneletti, D. (2010). Identifying priority areas for Forest Landscape Restoration in Chiapas (Mexico): An operational approach combining ecological and socioeconomic criteria. Landscape and Urban Planning, 94(1), 20–30.

Ortega, U., Ametzaga-Arregi, I., Sertutxa, U., & Peña, L. (2023). Identifying a green infrastructure to prioritise areas for restoration to enhance the landscape connectivity and the provision of ecosystem services. Landscape Ecology, 38(12), 3751–3765.

Ovaskainen, O. (2003). Habitat destruction, habitat restoration and eigenvector–eigenvalue relations. Mathematical biosciences, 181(2), 165–176.

Ovaskainen, O., & Hanski, I. (2003). How much does an individual habitat fragment contribute to metapopulation dynamics and persistence?. Theoretical population biology, 64(4), 481–495.

R Core Team. (2024). R: A language and environment for statistical computing (Version 4.4.1). R Foundation for Statistical Computing, Vienna, Austria. https://www.R-project.org/

Ranius, T., Martikainen, P., & Kouki, J. (2011). Colonisation of ephemeral forest habitats by specialised species: Beetles and bugs associated with recently dead aspen wood. Biodiversity and Conservation, 20(13), 2903–2915. 10.1007/s10531-011-0124-y

Saerens, M., Achbany, Y., Fouss, F., & Yen, L. (2009). Randomized shortest-path problems: Two related models. Neural Computation, 21(8), 2363–2404. 10.1162/neco.2009.11-07-643

Saura, S., & Rubio, L. (2010). A common currency for the different ways in which patches and links can contribute to habitat availability and connectivity in the landscape. Ecography, 33(3), 523–537.

Saura, S., Estreguil, C., Mouton, C., & Rodríguez-Freire, M. (2011). Network analysis to assess landscape connectivity trends: application to European forests (1990–2000). Ecological Indicators, 11(2), 407–416.

Saura, S. & J. Torné. 2009. Conefor Sensinode 2.2: a software package for quantifying the importance of habitat patches for landscape connectivity. Environmental Modelling & Software 24: 135–139.

Saura, S., & Pascual-Hortal, L. (2007). A new habitat availability index to integrate connectivity in landscape conservation planning: Comparison with existing indices and application to a case study. Landscape and Urban Planning, 83(2–3), 91–103. 10.1016/j.landurbplan.2007.03.005

Siitonen, J. (2001). Forest management, coarse woody debris and saproxylic organisms: Fennoscandian boreal forests as an example. Ecological Bulletins, 49, 11–41.

Strassburg, B. B., Iribarrem, A., Beyer, H. L., Cordeiro, C. L., Crouzeilles, R., Jakovac, C. C., … & Visconti, P. (2020). Global priority areas for ecosystem restoration. Nature, 586(7831), 724–729.

Suding, K. (2011). Toward an Era of Restoration in Ecology: Successes, Failures, and Opportunities Ahead. Annual Review of Ecology, Evolution, and Systematics, 42, 465–487. 10.1146/annurev-ecolsys-102710-145115

Sverdrup-Thygeson, A., Gustafsson, L., & Kouki, J. (2014). Spatial and temporal scales relevant for conservation of dead-wood associated species: current status and perspectives. Biodiversity and conservation, 23(3), 513–535.

Tambosi, L. R., Martensen, A. C., Ribeiro, M. C., & Metzger, J. P. (2014). A framework to optimize biodiversity restoration efforts based on habitat amount and landscape connectivity. Restoration ecology, 22(2), 169–177.

Tikkanen, O.-P., Martikainen, P., Hyvärinen, E., Junninen, K., & Kouki, J. (2006). Red-listed boreal forest species of Finland: Associations with forest structure, tree species, and decaying wood. Annales Zoologici Fennici, 43, 373–383.

United Nations Environment Programme, & Food and Agriculture Organization of the United Nations. (2021). Becoming #GenerationRestoration: Ecosystem restoration for people, nature and climate. United Nations Environment Programme.

Van Moorter, B., Kivimäki, I., Panzacchi, M., & Saerens, M. (2021). Defining and quantifying effective connectivity of landscapes for species’ movements. Ecography, 44(6), 870–884. 10.1111/ecog.05351

Van Moorter, B., Kivimäki, I., Panzacchi, M., Saura, S., Brandão Niebuhr, B., Strand, O., & Saerens, M. (2023a). Habitat functionality: Integrating environmental and geographic space in niche modeling for conservation planning. Ecology, 104(7), e4105.

Van Moorter, B., Kivimäki, I., Noack, A., Devooght, R., Panzacchi, M., Hall, K. R., … & Saerens, M. (2023b). Accelerating advances in landscape connectivity modelling with the ConScape library. Methods in Ecology and Evolution, 14(1), 133–145.

Van Moorter, B., Kivimäki, I., Panzacchi, M., Schouten, R., Wuyts, B., Niebuhr, B. B., … & Boussange, V. (2026). A unified framework for prioritizing habitat and connectivity conservation through analytical sensitivity. bioRxiv, 2026–01.

Wang, X., Svensson, J., Jonsson, B. G., Singh, N. J., Bubnicki, J. W., Lopéz-Peinado, A., … & Ardö, J. (2025). Where to restore: Connectivity forest for spatial prioritization in forest landscape restoration. Iscience, 28(9).

