## appendices for "Where conservation and restoration matter most: sensitivity-based prioritization of connected forest"

**Appendix S1. Parameterization of habitat permeability**

**Appendix S2: Rescaling ecological distance to meter-equivalent units and ecological proximity**

### Appendix S1. Parameterization of habitat permeability

Habitat permeability $P_{i,g}$ was modelled as a function of baseline matrix permeability, forest condition, and correspondence between local tree-species composition and the focal ecological group. For spatial unit *i* and ecological group $g$,

$$P_{i,g} = B + (1 - B)\tilde{T}_{i,g}C_i,$$

where *B* is baseline permeability, $C_i$ is forest condition, and $\tilde{T}_{i,g}$ is a transformed measure of correspondence between the cell and forest type $g$.

We set baseline permeability to

$$B = 0.25,$$

so that non-forest cells remained permeable but imposed substantially greater resistance to movement than high-condition forest. Thus, cells outside the forest mask were assigned ($P_{i,g} = 0.25$).

Within forest, the tree-species proportion $T_{i,g}$ was transformed as

$$\tilde{T}_{i,g} = \max\left(0.01, T_{i,g}\right)^{1/3}.$$

The cube-root transformation reduced differences among forest types relative to their effect on habitat quality. This reflects the assumption that forest dominated by a non-focal tree type may provide little or no habitat for a focal ecological group while still functioning as relatively favourable movement habitat compared with non-forest matrix. The lower bound of 0.01 ensured that forest cells retained a small forest-related permeability contribution even where the focal forest type was absent.

Permeability therefore approached its maximum where forest condition was high and tree-species composition closely matched the focal ecological group. To avoid values of exactly one in subsequent movement calculations, permeability values were capped at 0.99.

This parameterization was intended to represent broad contrasts between habitat and movement functions rather than species-specific movement responses. In particular, tree-species composition strongly constrained habitat quality in Equation 1, whereas its influence on permeability was deliberately attenuated through the transformation of $T_{i,g}$. Consequently, cells could contribute little or no habitat to a focal ecological group while still facilitating movement through the landscape.

### Appendix S2: Rescaling ecological distance to meter-equivalent units and ecological proximity

The expected cost calculation initially produces an accumulated movement cost between locations *s* and *t*, expressed in abstract cost units. To place this distance on an interpretable spatial scale, we derived a conversion between expected cost and geographic distance under baseline matrix permeability. We then used the resulting meter-equivalent ecological distance to define ecological proximity through the exponential distance-decay function used in the connectivity model.

We constructed a 400 × 400-cell raster with uniform permeability equal to the baseline value ($B = 0.25$). From the centre cell, we calculated both expected cost distance and Euclidean distance to all other cells. Euclidean distance showed an approximately linear relationship with cost distance (Fig. S1). The linear scaling relationship had a slope of 33 m per cost-distance unit, which was used to express ecological distance in meter-equivalent units:

$$d_{st} = 33c_{st},$$

where $c_{st}$ is ecological distance in cost units and $d_{st}$ is ecological distance in meter-equivalent units.

This transformation provides a geographic reference for movement cost rather than converting ecological distance into Euclidean distance. In heterogeneous landscapes, $d_{st}$ may be shorter or longer than the Euclidean separation between two locations: movement through high-permeability habitat reduces expected cost, whereas movement through low-permeability habitat increases it.

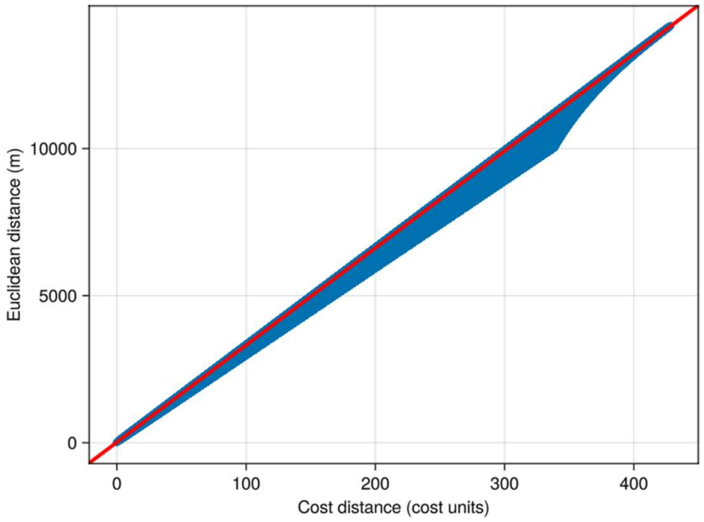


*Figure S1*. Relationship between accumulated movement cost and Euclidean distance under baseline matrix permeability. Cost distance and Euclidean distance were calculated from the centre of a 400 × 400-cell raster in which all cells were assigned the baseline permeability (B=0.25). The red line shows the linear scaling relationship used to convert ecological cost distance to meter-equivalent units; its slope of 33 m per cost-distance unit was used as the conversion factor.

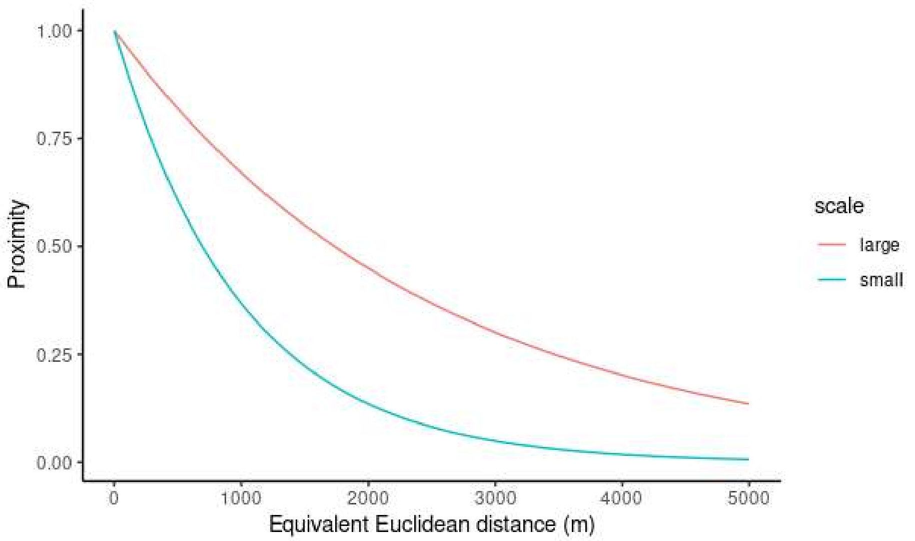


*Figure S2.* Relationship between ecological proximity and meter-equivalent ecological distance for the two movement-scale scenarios. Proximity was calculated using the exponential distance-decay kernel, $e^{-\alpha d_{st}}$, used in the connectivity model. The small-scale scenario declines more rapidly with distance ($1/\alpha = 1000m^{-1}$), representing more localized dispersal, whereas the large-scale scenario retains stronger connections over longer distances ($1/\alpha = 2500m^{-1}$).

The ecological distance $d_{st}$ was then used directly in the exponential connectivity kernel, $k_{st} = \exp(-\alpha d_{st})$, so that the movement-scale parameter $\alpha$ is expressed in the same meter-equivalent units (Fig. S2). We used two movement-scale scenarios ($1/\alpha \in \{1000; 2500\} m^{-1}$): the small-scale scenario representing more localized dispersal, whereas the large-scale scenario retains stronger connections over longer distances.
